# Elemental Accumulation in Kernels of the Maize Nested Association Mapping Panel Reveals Signals of Gene by Environment Interactions

**DOI:** 10.1101/164962

**Authors:** Greg Ziegler, Philip J. Kear, Di Wu, Catherine Ziyomo, Alexander E. Lipka, Michael Gore, Owen Hoekenga, Ivan Baxter

## Abstract

Elemental accumulation in seeds is the product of a combination of environment and a wide variety of genetically controlled physiological processes. We measured the kernel elemental composition of the Nested Association Mapping (NAM) of maize (*Zea mays* L.) grown in 4 different environments. Analysis of variance revealed strong effects of genotype, environment and genotype by environment interactions. Using Joint-linkage mapping on a set of 7000 markers we identified 354 quantitative trait loci (QTL) across 20 elements, four environments and a combination of the environments. Leveraging 20 M SNPs derived from genome resequencing on the parents of the population, genome-wide association mapping studies (GWAS) detected 8573 loci. While most of the GWAS SNPs were located near genes not previously implicated in elemental regulation, several SNPs were located next to orthologs of well-characterized elemental regulation genes.

## Introduction

Mineral elements are essential for all organisms to function, so over evolutionary time, all plants have developed and refined of processes to control their uptake, distribution and storage. To optimize fitness, organisms need to maximize uptake of limiting essential nutrients while reducing the uptake or sequestering chemically similar undesirable elements. This process is complicated by the wide variability of elemental availability, a function of parameters such as concentration, soil moisture, pH, cation-exchange-capacity, organic matter, and fertilizer applications [1]. The processes that respond to this variation in the environment to enable elemental uptake are poorly understood at the physiological and genetic level.

From its center of origin in Mexico, maize (Zea mays L.) has successfully adapted to many agro-ecological and environmental conditions due to its extensive genetic diversity [2]. Maize is now one of the most prevalent cereal crops in the world, with global production surpassing 1 billion tonnes in 2013 [3]. With a rich history as a genetic model, the maize community has created powerful genetic populations to use the genetic diversity to dissect the genetic control of many different traits. One such population is the maize nested association panel, derived from crossing 25 diverse parents to a single hub genotype and used to map loci for diverse traits such as flowering time, carotenoid concentration, and heat tolerance [4,5].

Ionomics, high throughput elemental profiling, has been a useful approach to understand elemental accumulation in both wild and crop plants. In addition to its use as a food source, the maize kernel is a defined step in the plants’ development and provides a highly heritable gauge of the genetic and environmental influences on elemental acquisition and homeostasis [6]. In this study, we have analyzed the elemental composition of kernels of the Maize NAM population grown in four diverse environments to identify the genetic loci underlying the ionome.

### Methods

The Maize Diversity project self pollinated ears of the Maize NAM in 2006 in North Carolina, New York, Puerto Rico and Florida as described in Buckler 2009 [4], and Brown [8]. Individual kernels from these growouts were analyzed on the ionomics pipeline as described in Ziegler et al.

Initially we analyzed the grow-outs in separate experimental batches containing only a single grow-out, but for later runs we mixed samples from separate locations, allowing us to remove batch effects from the analysis of environmental variation. All locations analyzed had some samples mixed in a batch with samples from other locations. The ratio between samples in these mixed runs was used to determine the true elemental ratio across locations. The elemental analysis encompassed over 100 runs of the ICP-MS and 50,000 total samples. Of the 18,000 packets analyzed, over 16,000 had two or more kernels analyzed.

### Outlier removal

Analytical outliers were removed from single-seed measurements using a method described in Davies and Gather [9]. Values were considered an outlier and removed from further analysis if the median absolute deviation (MAD) calculated based on the location and population each seed belongs to was greater than 10.

After outlier removal, an aggregate value for each analyte was calculated by taking the median value of the single-seed measurements for each line.

### Transformation

To meet the normality assumption required for further statistical analyses, the Box-Cox power transformation [10] was used to determine an appropriate transformation for each phenotype. Only the boron, aluminum, and cobalt phenotypes required a transformation.

### Calculation of BLUPs

Best linear unbiased predictors (BLUPs) of each ionomics compound for each line were predicted from a mixed model analysis across environments per Holland et al. [11] in ASReml version 3.0:

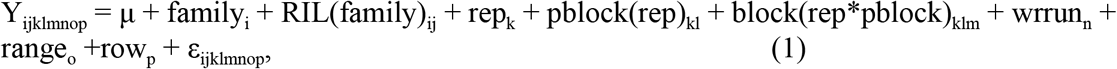

where Y_ijklmnop_ represents the phenotypic value of the p^th^ row of the o^th^ range of the n^th^ weighing robot run of the m^th^ block of the i^th^ pblock of the k^th^ rep of the j^th^ RIL of the i^th^ family, μ is the grand mean, family_i_ is the effect of the i^th^ family, RIL(family)_ij_ is the effect of the j^th^ RIL within the i^th^ family, repk is the effect of the kth rep, pblock(rep)_kl_ is the effect of the l^th^ pblock within the k^th^ rep, block(rep*pblock)_klm_ is the effect of the mth block within the l^th^ pblock and the the k^th^ rep, wrrun_n_ is the effect of the n^th^ weighing robot run, range_o_ is the effect of the o^th^ range, row_p_ is the effect of the p^th^ row, and and ε_ijklmnop_ represents a random error term. A first-order autoregressive (AR1× AR1) correlation structure was used to account for spatial variation among the rows and ranges. All terms except for family_i_ and RIL(family)_ij_ are random.

Likelihood ratio tests (testing H_0_: model term is not significant) were conducted to remove all terms from the model that were not significant at α = 0.05 [12]. The final model was then used to estimate BLUPs for each line and to estimate variance components.

### Joint-Linkage Analysis

Joint-linkage analysis was run using the StepwiseJointLinkagePlugin in TASSEL version 3.0 [13] with ~7,000 SNPs obtained by genotyping-by-sequencing from Elshire et al. 2011 [14]. To determine an empirical entry/exit p-value thresholds that control the type I error rate at α = 0.05, 1000 permutation tests were performed, in which the BLUP phenotype data were randomly shuffled within each NAM family before Joint Linkage analysis [15].

### Genome-wide Association

Stepwise forward regression genome-wide association was performed using the NamGwasPlugin in TASSEL version 4.0 [13] and is based upon previous NAM-GWAS experiments [16–18]. Briefly, genome-wide association was on a chromosome-by-chromosome basis. To account for variance explained by QTL on other chromosomes, the phenotypes used were the residuals from each chromosome calculated from the joint-linkage model fit with all significant joint-linkage QTL except those on the given chromosome. Association analysis for each trait was performed 100 times by randomly sampling, without replacement, 80% of the lines from each population.

The input SNP dataset was 28.9 million SNPs obtained from Maize Hapmap1 [19], Maize Hapmap2 [20] and an additional ~800,000 putative copy-number variants (CNVs) from analysis of read depth counts in Hapmap2 [16,20]. These ~30 million markers were projected onto the entire 5,000 line NAM population using low-density markers obtained through genotyping-by-sequencing [14]. A cutoff p-value of <1e-6 was used for inclusion in the final model. SNP associations were considered significant if selected in more than 5 of the 100 models (Resample model inclusion probability (RMIP) ≥ 0.05, [21]).

Data availability: All ionomic data, JL and GWAS results are available on the www.ionomicshub.org Data exchange. (Uploaded, 3/31/2016)

### Results & Discussion

#### Elemental profiling of the nested association mapping panel kernels across four locations

We conducted an ionomics analysis of the publicly available nested association-mapping (NAM) panel across four locations chosen for their widely differing environments (Table 1). The NAM panel combines high levels of genetic diversity while providing sufficient power to detect QTL. NAM is composed of 25 F6 generation recombinant inbred line (RIL) families, each with ~200 members, that all share B73 as a common parent [22,23]. A total of >50,000 kernels were analyzed for aluminum (Al), arsenic (As), boron (B), calcium (Ca), cadmium (Cd), cobalt (Co), copper (Cu), iron (Fe), potassium (K), magnesium (Mg), manganese (Mn), molybdenum (Mo), sodium (Na), nickel (Ni), phosphorus (P), rubidium (Rb), sulfur (S), selenium (Se), strontium (Sr) and zinc (Zn).

**Table 1.** Site soil type and chemistry of NAM population planting locations in 2006.

| Loc | Climat<br>e | pH | Soil Type | Substrate | Agronomy | Coordinates** |
| --- | --- | --- | --- | --- | --- | --- |
| FL | Subtro<br>p | 6.7 | Sand-clay<br>loam | Limestone<br>dolomite | High OM,<br>fertilized | 25°30'14.87" N<br>80°30'18.77" W |
| PR | Trop | 7.5 | San Anton<br>clay loam | Alluvial<br>deposit | Wet / fertile<br>High CEC | 18° 0'11.17" N<br>66°31'8.43" W |
| NC | Temp | 4.5 | Norfolk<br>loamy sand | Loamy<br>marine dep | Fertilized | 35°40'10.05" N<br>78°30'16.67" W |
| NY | Temp | 6.7 | Lima silt<br>loam | Limestone<br>calcareous | Fertilized | 42°43'31.18" N<br>76°39'5.10" W |

\*\* Coordinates provided by © Google 2016 from Google Earth Soil data provided by Natural Resource Conservation Service [7]

With a naive model including only location, and comparing >3500 packets from each location, all elements showed a significant effect (p<0.0001) of location. The difference between the median concentrations across environments varied from Mg (2%) to Sr (330%) (excluding Co which had a large difference between environments which were likely due to analytical issues owing to its extremely low concentration in seeds)(Table 2 and Figure 1). Soil properties such as underlying substrate and pH, as well as agronomic practices, are predicted to have strong effects on elemental accumulation. However, even though the NC soil is 2.2 pH units lower than all the other locations it didn't have the highest levels of Fe or other acid soluble elements. This suggests that the simple soil parameters are not the major drivers of differences in elemental accumulation data collected between locations.

**Table 2.** Percent difference of the median concentrations for each element between the highest and lowest grow-out.

| Element | Difference (%) |
| --- | --- |
| Magnesium | 2 |
| Potassium | 3 |
| Phosphorus | 6 |
| Sulfur | 8 |
| Boron | 10 |
| Arsenic | 10 |
| Manganese | 12 |
| Iron | 18 |
| Aluminum | 21 |
| Selenium | 30 |
| Zinc | 38 |
| Calcium | 38 |
| Copper | 56 |
| Molybdenum | 57 |
| Sodium | 77 |
| Cadmium | 220 |
| Nickel | 255 |
| Rubidium | 263 |
| Strontium | 330 |

**Figure 1.**
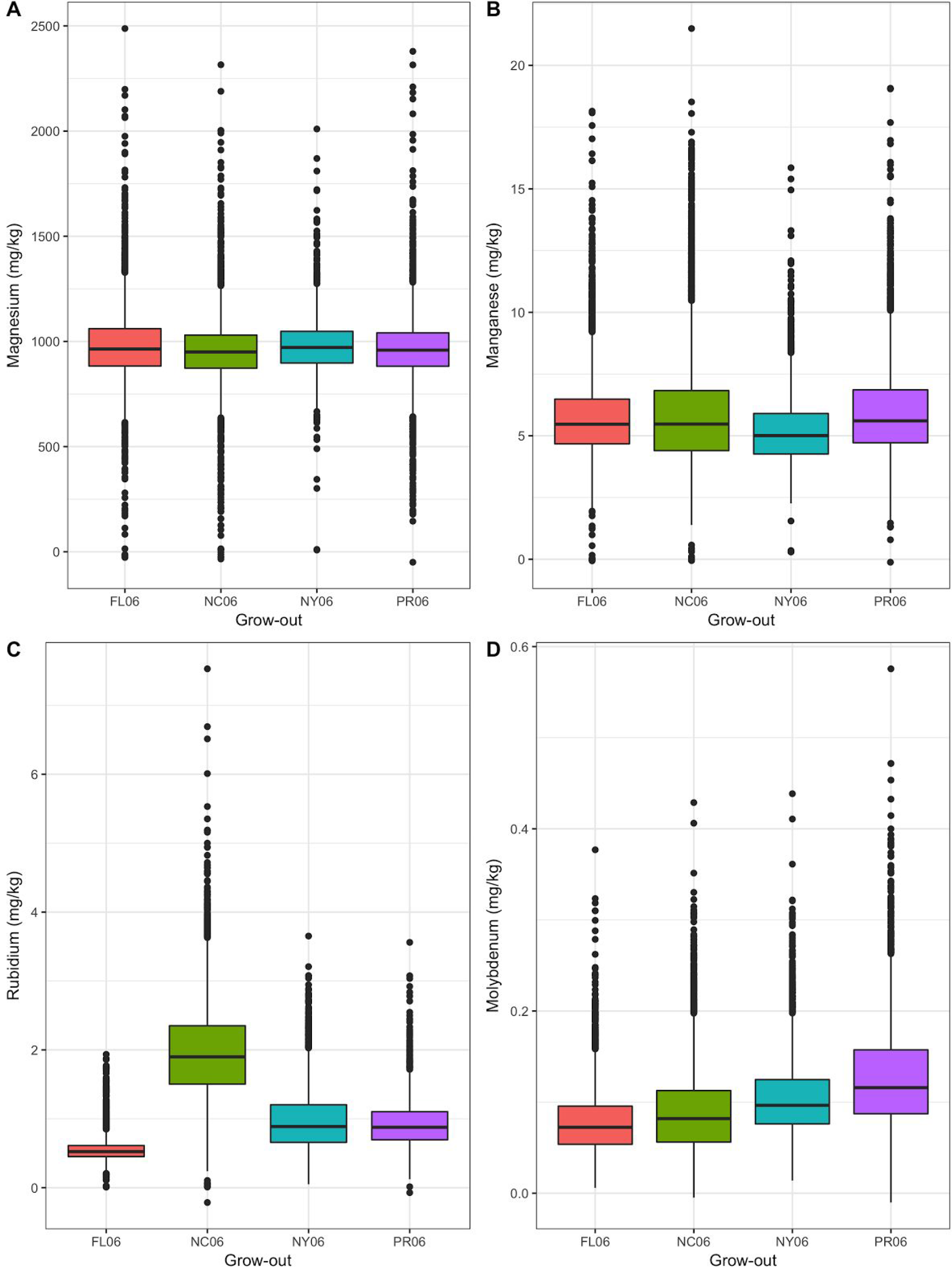
Concentrations of magnesium, manganese, rubidium, and molybdenum in NAM RIL kernels, a subset of twenty elements analyzed, at four locations. Concentrations (ppm) magnesium, manganese, rubidium, and molybdenum in Florida (FL06; red), North Carolina (NC06; green), New York (NY06; blue) and Puerto Rico (PR06; purple) in 2006 from a sample size of approximately three maize kernels for each of ~3500 lines in each location analyzed using ICP-MS. The median (solid black horizontal line), 25%-75% quartile (colored boxes for location), higher/lower extreme concentration (solid black vertical line) and outliers (individual points).

## Heritability

In order to partition the variance and estimate genotype values, we calculated BLUPs using the spatial checks in the field design and analytical checks. The nested design of the NAM population and the replicated field design across the four environments enabled a detailed statistical analysis of the sources of variation underlying the elemental traits. We performed linear mixed modeling of each element for each RIL across environments in ASReml. Significant terms were nested within a model for single and multiple environments. Models were fit separately for RIL families across environments, and each trait across environments. The final model was used to estimate best linear unbiased predictors (BLUPs) for each line and to estimate the components of phenotypic variance. The inclusion of the RIL family term, enables a robust calculation of V_GxE_ despite the single replicate plots of most lines in each location.

Hung *et al* [24] introduced a combined model for the analysis of *H^2^* and variances in the NAM population using harmonic means across multiple locations [24]. When applied to the elemental dataset, four elements had a high *H^2^* across NAM families (> 0.60; Cu, Fe, Mn and Mo), 6 elements were moderate (≥ 0.30 - 0.59; Al, Ca, K, Mg and S), and 10 were low (≤0.29; As, B, Cd, Co, Na, Ni, P, Rb, Se and Sr) (Figure 2). These results suggest that the elemental profile is highly heritable within environments but strongly influenced by the environment in a genotype dependent manner. The large amount of genetic by environment variation we observed is in sharp contrast to the amount found by Pfeiffer et al and X using similar methods to study height and flowering time in the same populations. However a more appropriate comparison would be to carry out the same analysis on the exact same plots for the different traits, allowing a side by side comparison.

**Figure 2:**
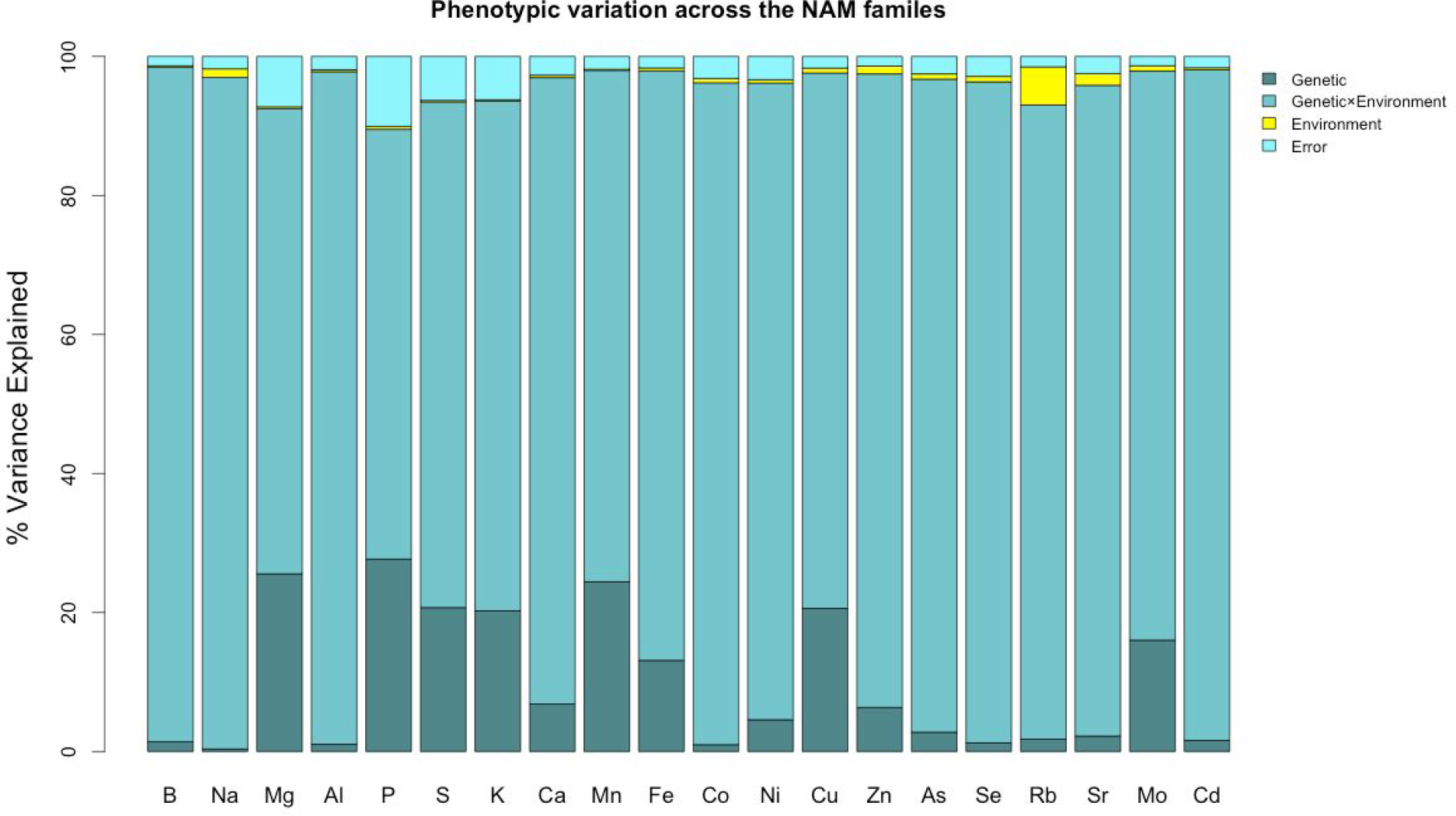
Partitioning of variance using the BLUP model.

### Joint Linkage Results

Stepwise joint-linkage regression was used for QTL detection across all 20 elements in the NAM population. QTL for each element were detected by using two methods: analyzing location data both individually and merged across locations using a best linear unbiased predictor (BLUP) model. In the four individual growouts, a total of 219 QTL were detected across 19 elements, with the exception of Se. Comparison of the individual location experiments revealed overlapping loci across multiple locations for a given element, clearly demonstrating that several QTL are derived from the same loci (Table 3).

**Table 3:**
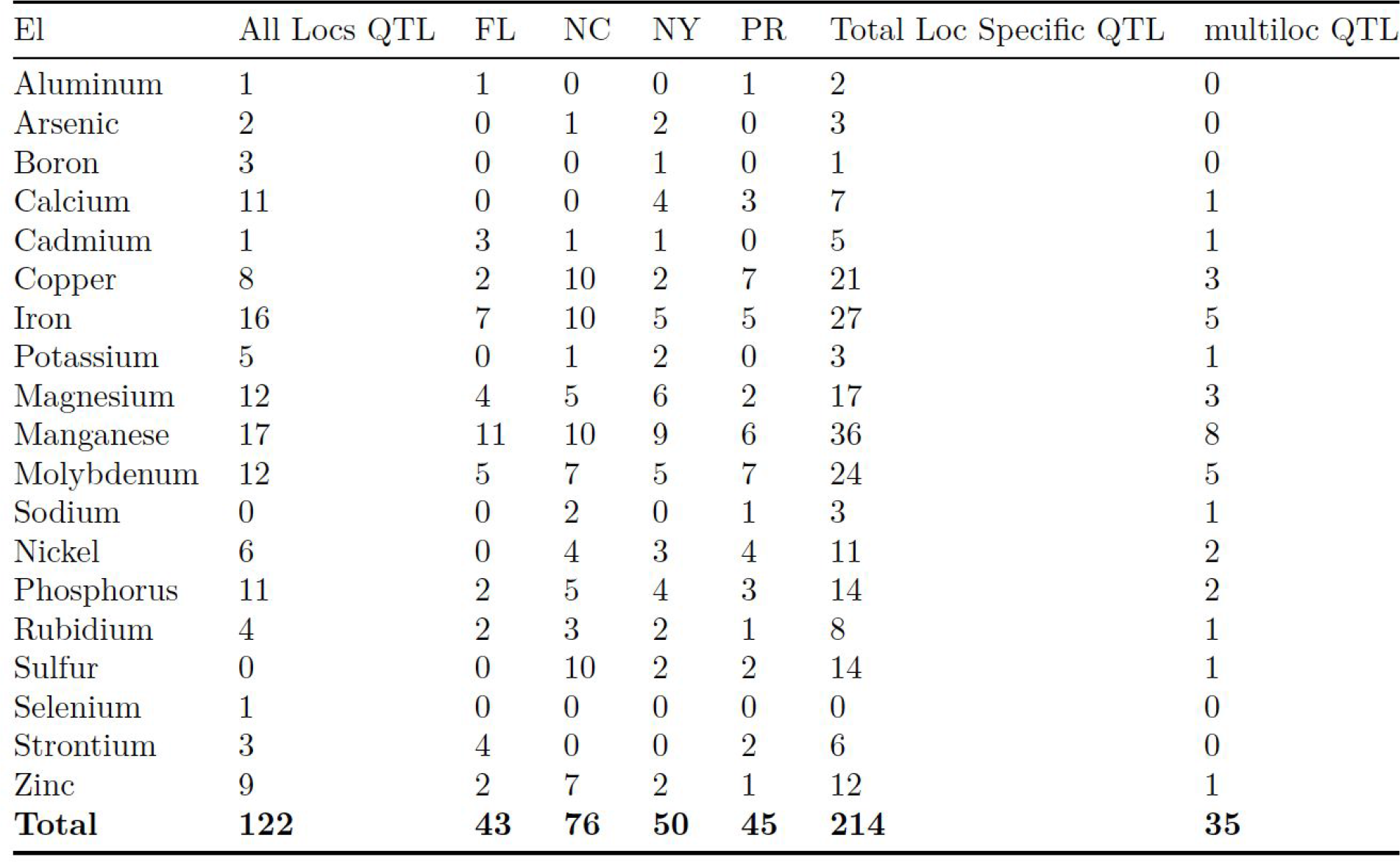
Summary of results for a Joint Linkage Analysis on four NAM grow-outs and a combined BLUP analysis. All locs QTL: Number of QTL detected from the all locations BLUP analysis; Total Loc specific QTL: Number of QTL detected for each element in the four locations; multi-loc QTL: Number of QTL whose 95% confidence interval overlaps between two or more locations.

### GWAS Results

The Joint-Linkage QTLs were used to account for loci on other chromosome as each chromosome was scanned for association with the 28M SNP markers that had been imputed on the full population. Using a cutoff of 5 iterations (out of 100) in the resampling approach, we identified a total of 8573 significant GWAS associations: 2923 in the all locations analysis and 5650 in the four locations specific datasets (Table 4). Counting overlap only as the identical SNP returned for the same element, we identified 29 loci that were found in multiple locations. Counting SNPs in LD with each other would result in many more SNPs being found in multiple locations.

**Table 4:** Summary of results from GWAS analysis of four NAM grow-outs and a combined location BLUP analysis. All locs QTL: Number of QTL detected from the all locations BLUP analysis; Total Loc specific QTL: Total number of QTL detected for each element in the four locations; multi-loc QTL: Number of instance where the exact same QTL was returned in two or more locations.

| El | All Locs QTL | FL | NC | NY | PR | Total Loc Specific QTL | multiloc QTL |
| --- | --- | --- | --- | --- | --- | --- | --- |
| Aluminum | 176 | 79 | 0 | 0 | 88 | 167 | 0 |
| Arsenic | 182 | 0 | 274 | 54 | 0 | 328 | 0 |
| Boron | 108 | 0 | 0 | 31 | 0 | 31 | 0 |
| Calcium | 105 | 0 | 0 | 53 | 55 | 108 | 0 |
| Cadmium | 630 | 72 | 115 | 45 | 0 | 232 | 1 |
| Copper | 165 | 143 | 102 | 89 | 97 | 431 | 4 |
| Iron | 171 | 130 | 112 | 61 | 83 | 386 | 4 |
| Potassium | 130 | 0 | 253 | 47 | 0 | 300 | 1 |
| Magnesium | 153 | 89 | 84 | 78 | 84 | 335 | 0 |
| Manganese | 168 | 115 | 111 | 94 | 110 | 430 | 8 |
| Molybdenum | 154 | 124 | 102 | 123 | 140 | 489 | 6 |
| Sodium | 0 | 0 | 320 | 0 | 96 | 416 | 1 |
| Nickel | 99 | 0 | 100 | 78 | 144 | 322 | 2 |
| Phosphorus | 123 | 84 | 91 | 73 | 68 | 316 | 1 |
| Rubidium | 135 | 118 | 91 | 71 | 151 | 431 | 0 |
| Sulfur | 0 | 0 | 91 | 70 | 91 | 252 | 0 |
| Selenium | 162 | 0 | 0 | 0 | 0 | 0 | 0 |
| Strontium | 113 | 91 | 0 | 0 | 95 | 186 | 1 |
| Zinc | 149 | 52 | 99 | 221 | 118 | 490 | 0 |
| **Total** | **2923** | **1097** | **1945** | **1188** | **1420** | **5650** | **29** |

### High Confidence Candidate Genes

Visual inspection of the genes proximal to strong GWAS peaks revealed several genes whose orthologs are known to have roles in elemental accumulation (Table 5). These candidates suggest that the ionomics approach can be used with the NAM to identify maize genes important for elemental accumulation in the kernel.

**Table 5.**
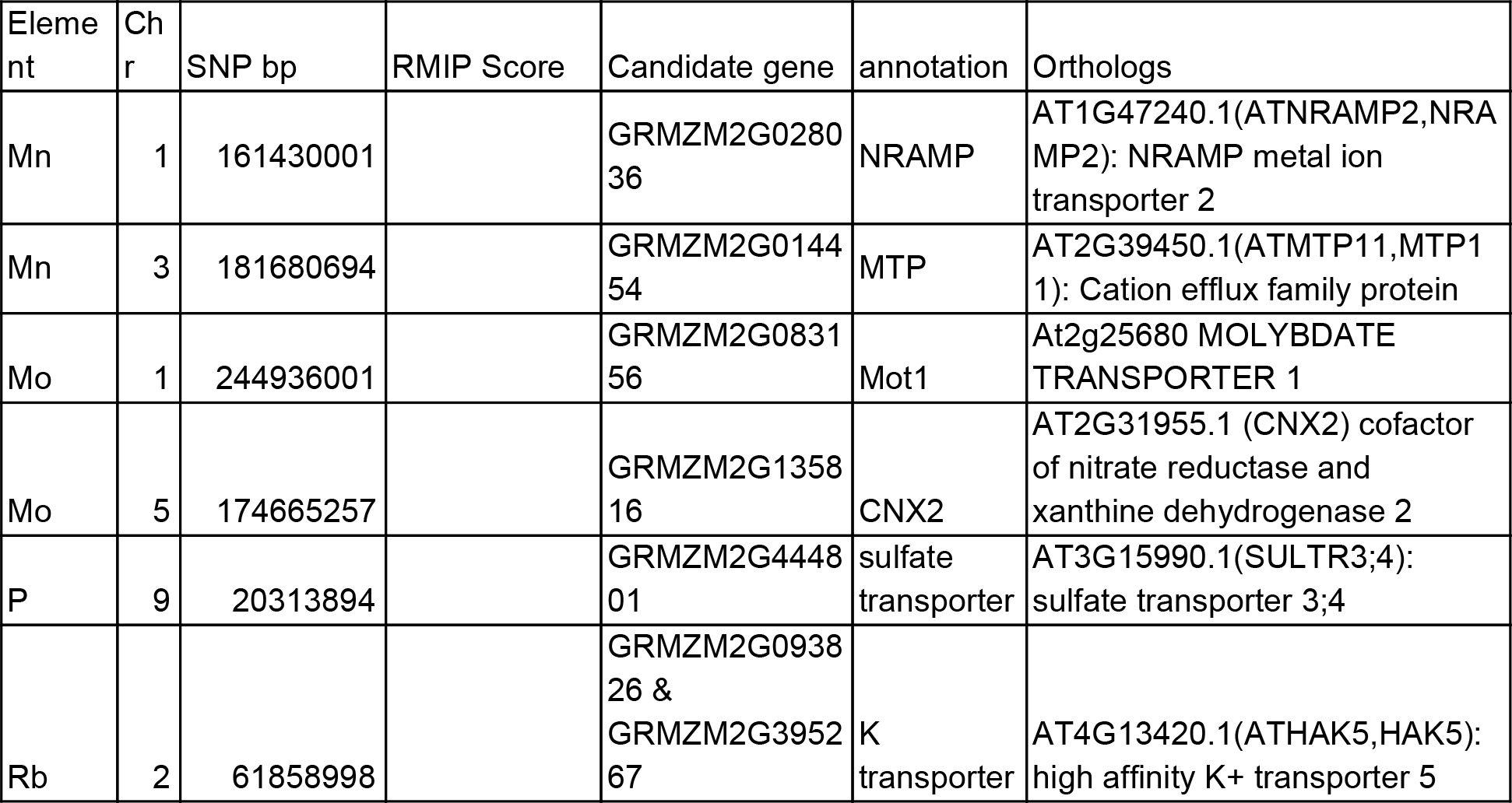
High Confidence genes identified under GWAS peaks

### Manganese

Manganese is an essential plant element that is a cofactor in many crucial metalloenzymes, such as those in photosystem II [25]. Various methods of manganese transport and homeostasis in plants have been characterized [26,27]. Exploring the GWAS results for a model that combined the data from the four grow-out locations reveals a strong association (RMIP=0.98) for a SNP on chromosome 1 (Figure 3). In three of the four grow-outs, this SNP is strongly associated with manganese accumulation. The gene directly under this SNP is a putative natural resistance-associated macrophage protein (NRAMP2) metal ion transporter (Figure 3). The second highest SNP for the combined location GWAS is on chromosome 3 with an RMIP of 0.83 (Fig 3). This SNP is significant in all four of the locations and it falls directly in a gene annotated as manganese transporter protein 11 (MTP11) homolog. MTP11 is a member of the cation diffusion facilitator (CDF) family and is known to have roles in known to have roles in manganese tolerance and homeostasis [28,29].

**Figure 3.**
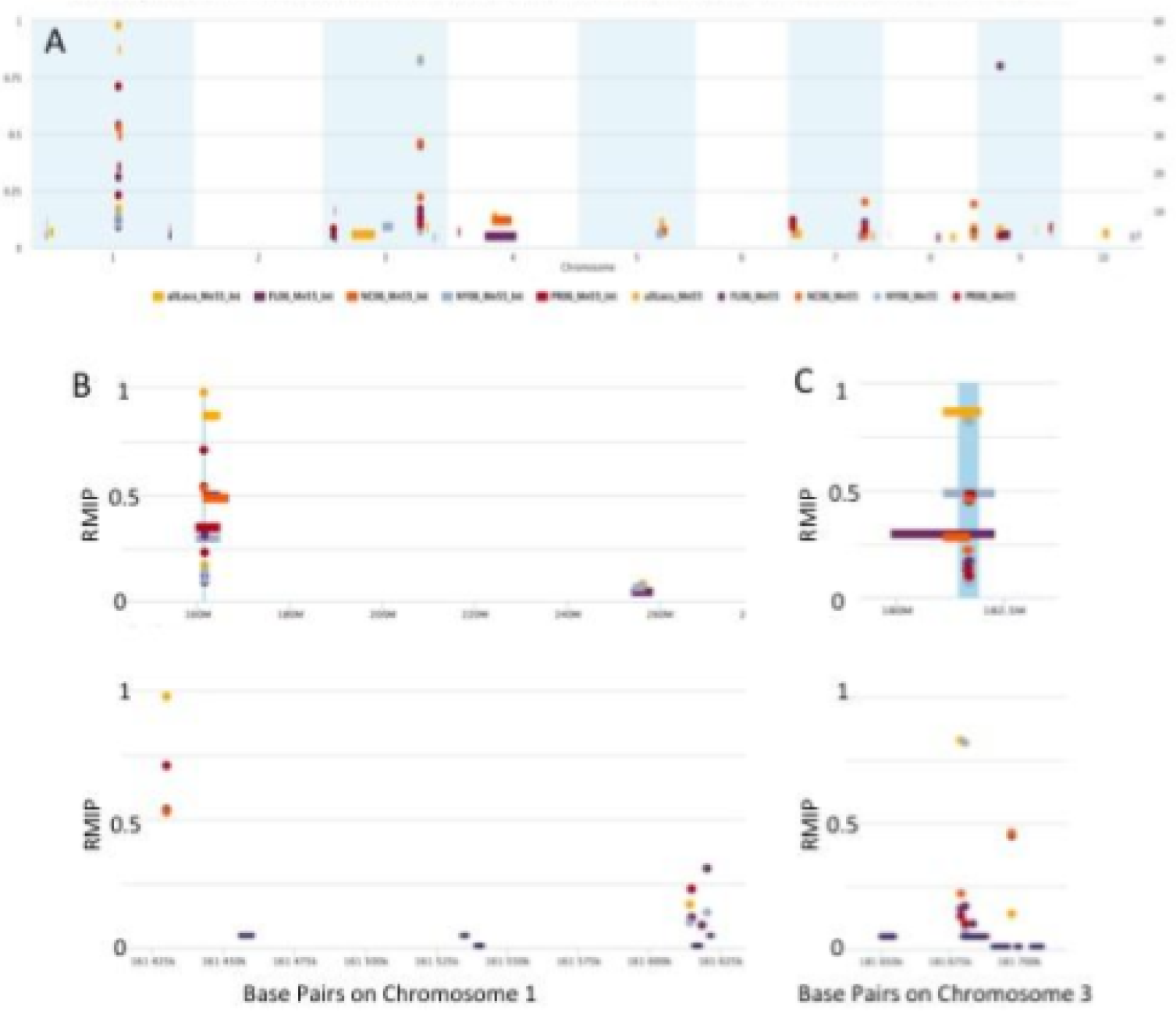
Multilocation hit for tow candidate genes contributing to managanese accumulation. A. Whole genome view showing GWAS and Joint Linkage hits. B. Chromosome level (top) and Gene level (bottom) zoom of chromosome 1 peak. C. Chromosome level (top) and Gene level (bottom) Zoom of chromosome 3 peak.

### Molybdenum

We identified 13 unique QTL for Mo in the joint linkage analysis, two of which were extremely strong (F-test of 32 and 18 with a experiment wide significance cutoff < 3) (Figure 4). Candidate genes for Mo accumulation include those from the biosynthetic pathway of the molybdopterin cofactor and Mo transporters with orthologs in Arabidopsis. A biosynthetic gene and the maize ortholog of the Arabidopsis Mot1 transporter are located under the top two joint linkage peaks and thus represent strong candidates for the likely quantitative trait genes (QTGs).

**Figure 4.**
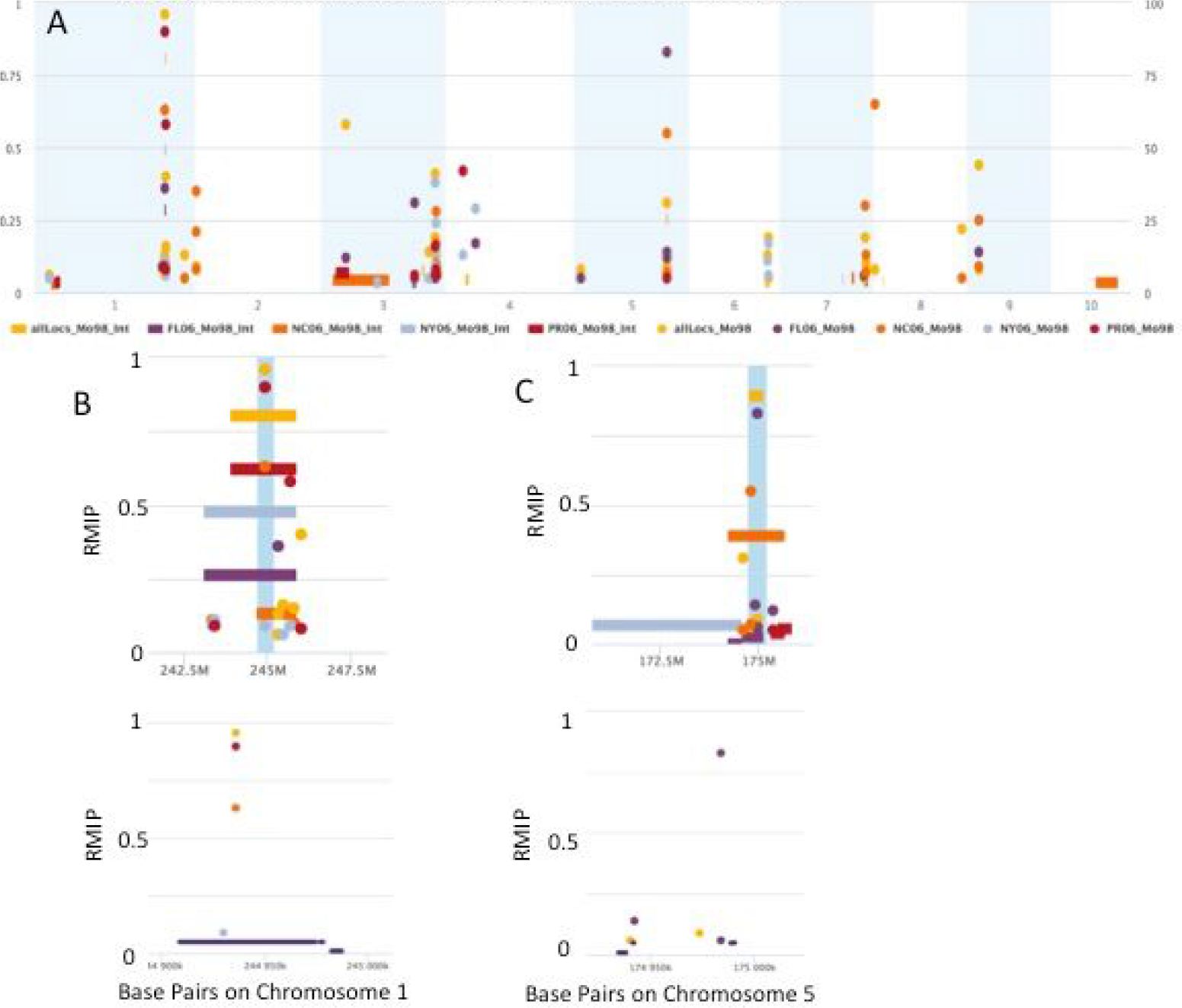
Manhattan plot showing Joint Linkage and GWAS results for molybdenum in 4 locations and 1 combined location. A. Whole genome view showing GWAS and Joint Linkage hits. B. Chromosome level (Top) and Gene Level (Bottom) zoom of peak over Mot1 on chromosome 1. C. Chromosome level (Top) and Gene Level (Bottom) zoom of peak over Mot1 on chromosome 5.

### Molybdenum Gene by Environment Interaction

The two top molybdenum loci showed a strong reciprocal gene by environment interaction (Figure 5). While both loci were significant in the joint linkage in all four environments (and the all locations), in FL, PR and NY, the Chr 1 loci had a substantially higher F statistic, while in NC the Chr. 5 loci was stronger than Chr 1. While we do not know what environmental parameters drove this difference, the growth environment is having a strong effect on the genetic cause of the phenotype.

**Figure 5.**
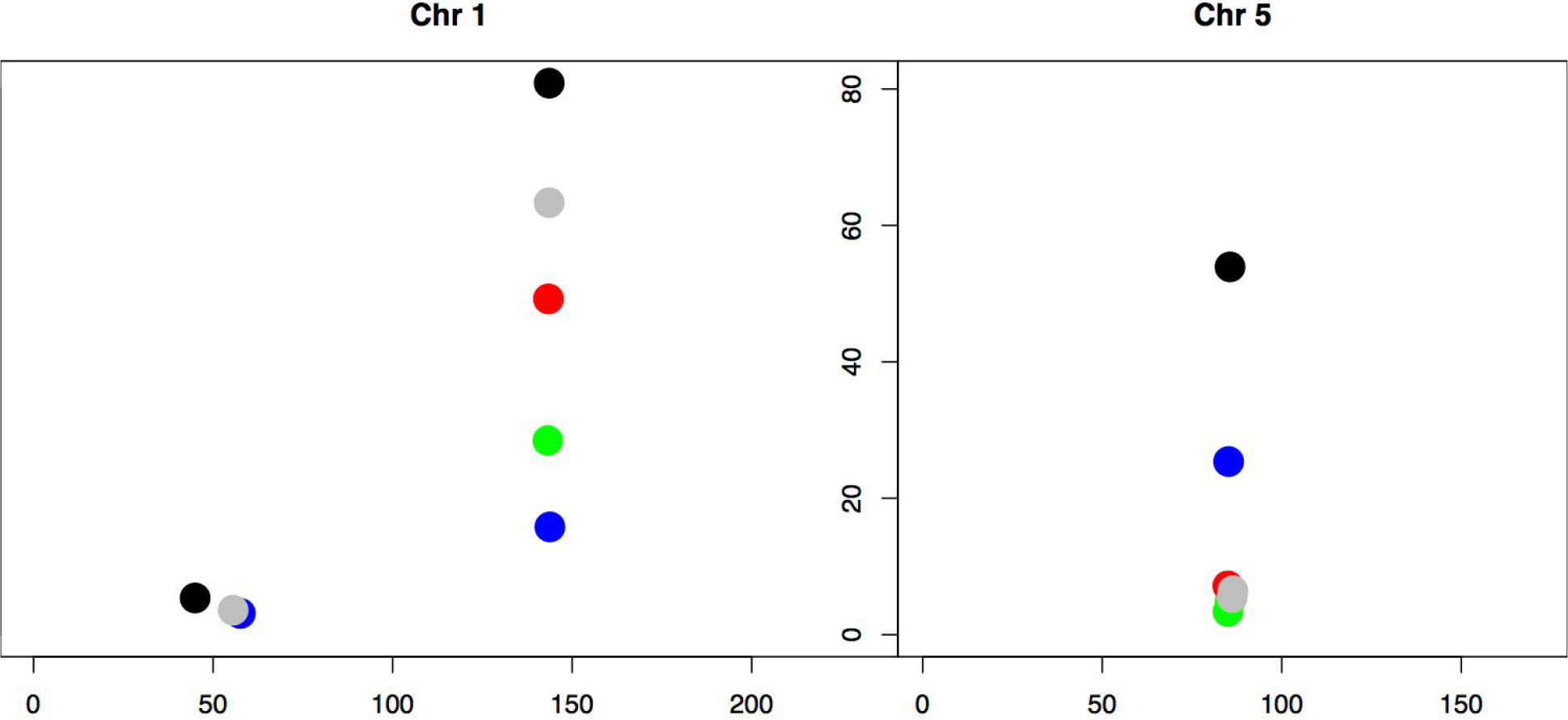
F-value of the two strongest Molybdenum loci in the four locations and the all locations model. Black (all locs), Green (FL), Blue (NC), Red (NY), and Grey (PR) denote the locations.

### Phosphorus

A phosphorus QTL on chromosome 9 was returned for 3 out of the 4 locations (NC, NY, and FL) (Figure 6). A likely candidate gene for this peak is GRMZM2G444801 which is an ortholog of *low-phytic acid-1* (*lpa1*) in barley. In barley, mutations in *lpa1* heavily influence seed phosphorus composition [30]. Interestingly, in Maize, *lpa1* mutants have been characterized as altering the balance of phytic acid to inorganic phosphorous, but not the total phosphorus content [31]. However, from our results, it appears that diversity in *lpa1* does have some phenotypic effect on total phosphorus content.

**Figure 6.**
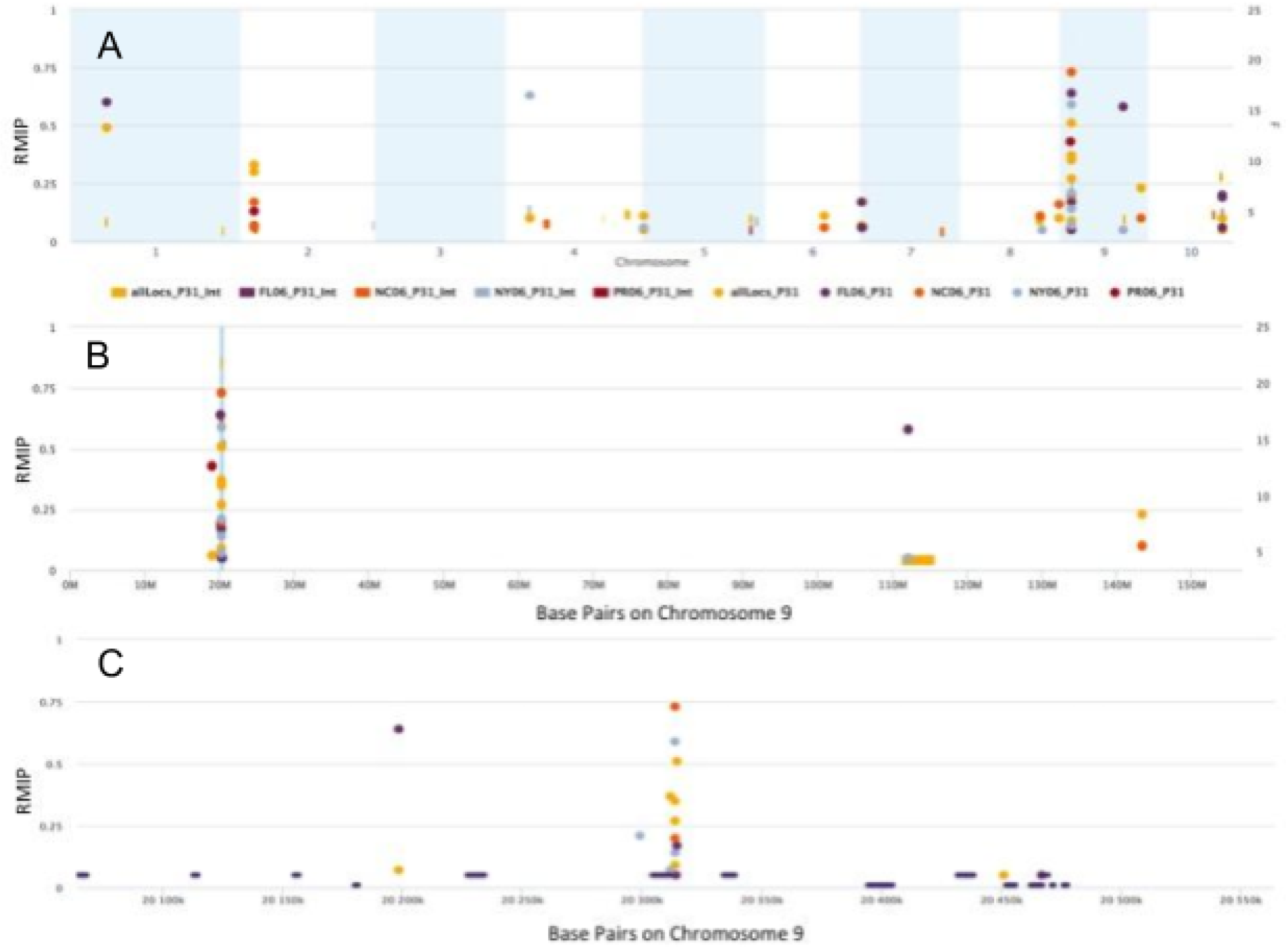
Manhattan plot of GWAS and Joint Linkage results for phosphorus. Displaying SNPs found in two or more locations. A. Genome-wide manhattan plot. B. Chromosome level zoom to peak on chromosome 9. C. Gene-level zoom to peak on chromosome 9.

### Rubidium

The most significant hit across the 4 locations tested for rubidium in both the joint linkage and GWAS analysis was a SNP on chromosome 2. This SNP was only returned as significant in the New York growout. Interestingly, the New York location did not have the highest median accumulation of rubidium (Figure 1). This rubidium hit is very close to two orthologs of an Arabidopsis high-affinity potassium transporter (Figure 7). In Arabidopsis, this gene has been shown to play a role in potassium acquisition under low-potassium conditions [32]. In sunflowers, it has been shown that there is a change in rubidium uptake kinetics depending on potassium availability [33]. The unique interplay between environmental factors and genetics found in the New York NAM grow-out resulted in a significant difference in rubidium accumulation in plants.

**Figure 7.**
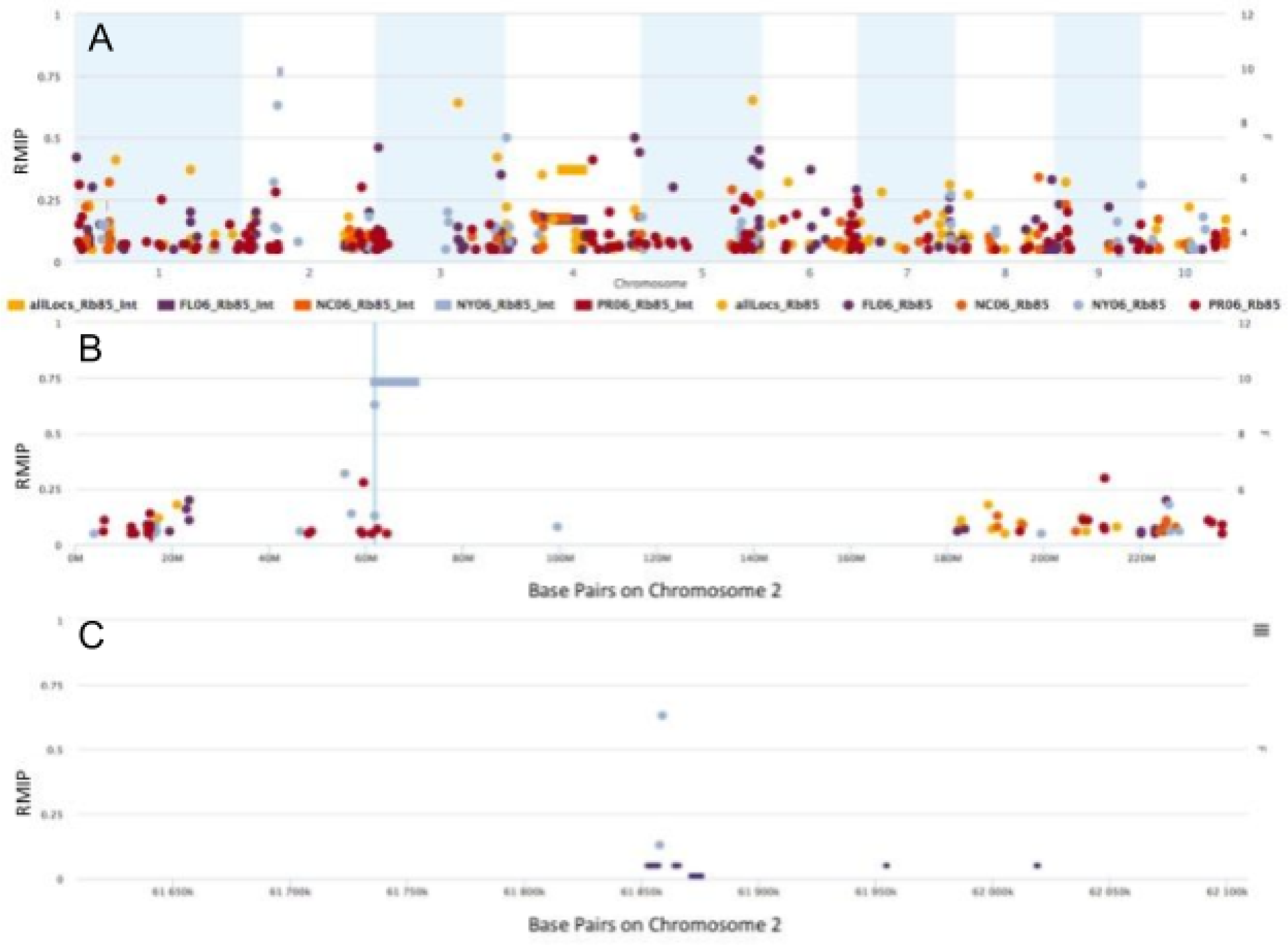
Manhattan plot of rubidium GWAS and Joint Linkage results. A. Whole genome manhattan plot.B. Chromosome-wide manhattan plot.C.Single location hit on chromosome 2 next to a potassium transporter.

### Conclusion

Here we have analyzed the elemental composition of over 50,000 kernels of the maize Nested association mapping population grown in four locations. We demonstrate that the elemental composition is heritable with a large genetic by environment interaction. Using a two step genetic mapping approach, we identified >300 loci controlling 19 of the 20 elements that we measured. Strong signal was observed around orthologs of known elemental accumulation genes. This data will be a rich resource for identification of the genes driving elemental accumulation in maize.

## Acknowledgements

The authors would like to thank the Maize Diversity project for creating and sharing the resources used in this project. They would also like to thank Jason Wallace and Margaret Smith for technical assistance and Cyverse for computational support. The research was funded by U.S. National Science Foundation, Plant Genome Research Program (grant #IOS 1126950 awarded to Ivan Baxter and Owen Hoekenga, IOS-0922493 (past) and IOS-1546657 (current) awarded to Michael Gore) and United States Department of Agriculture-Agricultural Research Service Intramural funds to Ivan Baxter. The funders had no role in study design, data collection and analysis, decision to publish, or preparation of the manuscript.

